# Circulating metabolites of *Borneolum syntheticum* (Bingpian) inhibit foam-cell formation in macrophages induced by oxidized low-density lipoprotein

**DOI:** 10.1101/2023.07.10.548303

**Authors:** Rong-rong He, Hui Li, Zi-xuan Chu, Feng-qing Wang, Fei-fei Du, Fang Xu, Jia-qi Wang, Olajide E. Olaleye, Ting Wang, Chen Cheng, Chuan Li

**Author notes:** Correspondence: Chen Cheng or Chuan Li. These authors contributed equally: Rong-rong He, Hui Li, Zi-xuan Chu.

## Abstract

Coronary heart disease is caused by the accumulation of atherosclerotic plaques which narrow the arteries over time. The plaques are formed by cholesterol deposits in the arterial intima and lead to the symptom of angina pectoris. *Borneolum syntheticum* (Bingpian) has been extensively used as a component in Chinese herbal medicines for cardiovascular diseases. This investigation aimed to examine Bingpian metabolism and its effects on anti-atherosclerotic activities. Major circulating Bingpian compounds were detected in human subjects who received a Bingpian-containing medicine. In vitro and rat studies were also conducted to facilitate the understanding of disposition factors that govern the systemic exposure to Bingpian compounds. Although Bingpian constituents, borneol (**1**) and isoborneol (**2**), are efficiently absorbed in the intestine, extensive hepatic first-pass glucuronidation, which is mediated predominantly by UGT2B7, coupled with MRP3 and MRP4-mediated efflux of the glucuronides into the blood, and oxidation, which is mediated by CYP2A6, CYP2B6, and CYP3A, result in the formation of metabolites borneol-2-*O*-glucuronide (**M1_G_**), isoborneol-2-*O*-glucuronide (**M2_G_**), and camphor (**3**) as the major circulating Bingpian compounds instead of the unchanged **1** and **2**. Glucuronides are predominantly eliminated through renal excretion, which involves both glomerular filtration and OAT3- and OAT4-mediated tubular secretion. Furthermore, **M1_G_**, **M2_G_**, and **3**, as well as **1** and **2**, displayed inhibitory effects on oxidized low-density lipoprotein-induced foam-cell formation in macrophages. The findings emphasized that the metabolites must be given priority in pharmacodynamic studies of Bingpian. Comprehensive integration of pharmacokinetic and pharmacodynamic studies facilitates understanding how Bingpian functions in the body to provide therapeutic benefits.

## INTRODUCTION

Coronary heart disease is a progressively narrowing of the arteries that occurs over time due to the gradual accumulation of atherosclerotic plaques. This condition, often accompanied by angina pectoris, is a significant public health concern worldwide [1, 2]. Current medical therapy for angina pectoris involves antianginal medications designed to reduce angina frequency and improve quality of life, as well as preventive medications intended to lower the risk of cardiovascular events [3]. In China, traditional herbal medicines are frequently employed for treatment and prevention of angina pectoris [4]. *Borneolum syntheticum*, also known as "Bingpian" or synthetic Bingpian in Chinese, is widely used as a component in many cardiovascular Chinese herbal medicines. The Pharmacopoeia of the People’s Republic of China (2020 edition) lists 36 cardiovascular herbal medications containing Bingpian [5].

Bingpian, used in Chinese traditional medicine, was originally a medicinal spice imported from Southeast Asia more than a millennium ago [6]. This type of Bingpian (*Borneolum* or natural Bingpian) is a crystal form of *d*-borneol obtained through distillation of the trunk of *Dryobalanops aromatica* Gaertn. F. (family *Dipterocarpaceae*). However, due to limited availability of the *D. aromatica* tree, synthetic Bingpian has become a commonly used alternative since 1950s. This synthetic Bingpian comprises a mixture of *dl*-borneol and *dl*-isoborneol (with borneol content of at least 55%) and is prepared through the esterification and hydrolysis of α-pinene from turpentine or the reduction of camphor. In 1980s, Chinese botanists discovered *Cinnamomum camphora* (L.) Presl (family *Lauraceae*) trees that had rich reserves of *d*-borneol within the country. Since then, natural Bingpian from domestic *C*. camphora trees has been used and is listed in the Chinese Pharmacopoeia. However, the use of natural Bingpian is only limited to four ophthalmic herbal medicines, whereas synthetic Bingpian is present in all cardiovascular herbal medicines listed in the Chinese Pharmacopoeia [5]. Several animal pharmacological studies suggested that synthetic Bingpian (referred to as ‘Bingpian’ thereafter) exhibits various properties such as vascular protection, anticoagulation, neuroprotection, sedation, anti-inflammation, antioxidation, and analgesia [7⍰15]. In addition, ex vivo and cell-based studies have demonstrated endothelium-independent vasorelaxant activities of *l*-borneol on rat thoracic aorta artery rings, inhibitory activities of *dl*-isoborneol on accumulation of lipids in macrophages, and anti-inflammatory and antioxidant activities of borneol on rat neurons harmed by oxygen-glucose deprivation followed by reperfusion [16⍰18]. However, some pharmacokinetic studies suggest low oral bioavailability of both borneol and isoborneol (11.9%–14.0% and 2.0%–4.9%, respectively) in rats administered with Bingpian products [19⍰21]. Given favorable membrane permeability and absorption-supporting solubility of the monoterpene alcohols, their low bioavailability is hypothesized to result from first-pass metabolism. Nonetheless, knowledge of the metabolism of Bingpian, particularly that in humans, is still restricted [22].

The aim of this investigation was to examine systemic exposure to and metabolism of Bingpian in human subjects who orally received compound Danshen tablet, a Bingpian-containing cardiovascular herbal medicine, and to evaluate anti-atherosclerotic activities of Bingpian-derived compounds that were prioritized by the pharmacokinetic study. Notably, the metabolites, rather than the parent monoterpene alcohols, were identified as the predominant circulating compounds of Bingpian and exhibited the ability to reduce lipid accumulation and foam-cell formation in macrophages, induced by oxidized low-density lipoprotein (ox-LDL). The investigation has important implications for enhancing pharmacological research on Bingpian or Bingpian-containing cardiovascular herbal medicines through the investigation of the pharmacokinetics-selected ‘right’ compounds.

## MATERIALS AND METHODS

### Materials and reagents

Bingpian, which contains 60.0% borneol and 39.3% isoborneol, was acquired from Ji’an Lvyuan Natural Perfume Oil Refinery (Ji’an, Jiangxi Province, China). Compound Danshen tablet, a Chinese herbal medicine used to treat stable angina pectoris, was manufactured by Yunnan Baiyao Group Co. (Kunming, Yunnan Province, China). It has been approved by the National Medical Products Administration with a drug ratification number of GuoYaoZhunZi-Z53021243. The tablet was prepared from a three-herb combination comprising *Salvia miltiorrhiza* roots (known as Danshen in Chinese), *Panax notoginseng* roots (Sanqi), and *Borneolum syntheticum* (Bingpian). Each tablet (320 mg) is standardized to contain ≥ 0.2 mg tanshinone IIA, ≥ 5.0 mg salvianolic acid B, and ≥ 6.0 mg ginsenosides (i.e., the total amount of ginsenosides Rg_1_, Rb_1_, and Re and notoginsenoside R_1_). The compound Danshen tablets utilized in the current human study were from lot ZKC1424. Each tablet of this lot contained 2.64 mg (17.1 μmol) borneol, 1.88 mg (12.2 μmol) isoborneol, and 0.01 mg (0.04 μmol) camphor.

*dl*-Borneol was obtained from Shanghai Aladdin Bio-Chem Technology (Shanghai, China); *dl*-isoborneol and naphthalene (used as internal standard for GC-MS analysis), from the National Institutes for Food and Drug Control (Beijing, China); and camphor and *l*-menthol-*O*-glucuronide (used as internal standard for LC-MS/MS analysis), from J&K Scientific Co. (Beijing, China). Alamethicin, 4-methylumbelliferone, 4-methylpyrazole, oestradiol 17β-D-glucuronide, para-aminohippuric acid, prostaglandin F2α, estrone-3-sulfate, nicotinamide adenine dinucleotide phosphate (NADPH), nicotinamide adenine dinucleotide (NAD), glucose-6-phosphate monosodium salt, glucose-6-phosphate dehydrogenase, uridine 511-diphospho-glucuronic acid (UDPGA), and *tris*-hydroxymethyl-aminomethane were obtained from Sigma-Aldrich (St. Louis, MO, USA); and midazolam, cinnamyl alcohol, glycyrrhizin, and atorvastatin calcium, from Shanghai Standard Technology (Shanghai, China). The purity of these compounds all exceeded 98%.

Pooled human liver microsomes, human intestine microsomes, human liver cytosol, and human intestine cytosol were obtained from Corning Gentest (Woburn, MA, USA). cDNA-expressed human cytochrome P450 (CYP) enzymes (CYP1A2, CYP2A6, CYP2B6, CYP2C8, CYP2C9, CYP2C19, CYP2D6, CYP2E1, CYP3A4, and CYP3A5) and human uridine 5’-diphosphoglucuronosyltransferase (UGT) enzymes (UGT1A1, UGT1A3, UGT1A6, UGT1A7, UGT1A8, UGT1A9, UGT1A10, UGT2B4, UGT2B7, UGT2B10, UGT2B15, and UGT2B17) were obtained from Corning Gentest. Inside-out membrane vesicles expressing human multidrug resistance-associated protein (MRP) 2, MRP3, MRP4, breast cancer resistance protein (BCRP), bile salt export pump (BSEP), and multidrug resistance 1 (MDR1) were obtained from GenoMembrane (Kanazawa, Ishikawa Prefecture, Japan). Expression plasmids of human organic anion transporter (OAT) 1, OAT2, OAT3, OAT4, human organic anion-transporting polypeptide (OATP) 1B1, OATP1B3, and OPATP2B1 were constructed by Invitrogen Life Technologies (Shanghai, China). Human embryonic kidney 293 (HEK-293) cells and murine macrophage RAW 264.7 cells were obtained from American Type Culture Collection (Manassas, VA, USA). Dulbecco’s modified Eagle’s medium (DMEM), minimal essential medium non-essential amino acids, and penicillin-streptomycin were obtained from Life Technologies (Grand Island, NY, USA); fetal bovine serum (FBS), from Corning Gentest; oxidized low-density lipoprotein (ox-LDL), 1,1’-dioctadecyl-3,3,3’,3’-tetramethyl-indocarbocyanine perchlorate-labeled ox-LDL (DiI-ox-LDL), and high-density lipoprotein (HDL), from Guangzhou Yiyuan Biotechnology (Guangzhou, Guangdong Province, China); Oil Red O stain kit and ELISA kits for total cholesterol and free cholesterol, from Applygen Technologies Inc. (Beijing, China); cell counting kit-8 (CCK-8), DAPI staining solution, bicinchoninic acid protein assay kit, and radio immunoprecipitation assay (RIPA) lysis buffer, from Beyotime Biotechnology (Shanghai, China); and 22-(7-Nitrobenz-2-oxa-1,3-diazol-4-yl-amino)-23,24-bisnor-5-cholen-3β-ol (22-NBD-cholesterol), from Thermo Fisher Scientific (Waltham, MA, USA). Sodium heparin and isoflurane were obtained from Sinopharm Chemical Reagent (Shanghai, China); and sodium taurocholate, from Solarbio Life Science (Beijing, China).

HPLC-grade methanol, acetonitrile, formic acid, and dimethyl sulfoxide (DMSO) were obtained from Sinopharm Chemical Reagent. Deionized water was purified using a Millipore Milli-Q Integral 3 cabinet water purifying system (Bedford, MA, USA).

### Human pharmacokinetic study

A single-center, open-label pharmacokinetic study of compound Danshen tablet was performed in healthy volunteers at the National Clinical Research Center of the Second Affiliated Hospital of Tianjin University of Traditional Chinese Medicine (Tianjin, China). The study protocol was reviewed and approved by an ethics committee of clinical investigation at the hospital. The study was registered on the Chinese Clinical Trials Registry (www.chictr.org.cn), with a registration number of ChiCTR-OPN-15006554 and performed in compliance with the Declaration of Helsinki. All volunteers provided written informed consent prior to enrollment.

Twelve participants, comprising six male and six female, were recruited for this study. Each participant ingested a single oral dose of 27 compound Danshen tablets on the first day of the study, which is thrice the recommended daily dose. Blood samples, approximately 3 mL each time, were drawn via an antecubital vein catheter, before and at 0.167, 0.3, 0.5, 1, 2, 4, 6, 9, 12, 16, 24, 29, 34, and 48 h following drug administration. Meanwhile, urine samples were collected before and at 0–6, 6–12, 12–24, 24–34, and 34–48 h after dosing. On the third day of the study, six participants (three male and three female) were selected from the 12 participants to receive the same dose of compound Danshen tablets for the following seven days. Blood samples were collected from the selected participants before and at 2 and 24 hours after daily dosing on days 3-8. The blood and urine sampling schedules on day 9 were identical to those on day 1. Heparinized blood samples were centrifuged at 3000 *g* for 10 min to obtain plasma fractions, while urine samples were not preserved. The plasma and urine samples were kept at -70°C until analyzed. Serum alanine aminotransferase, aspartate aminotransferase, total protein, albumin/globulin ratio, total bilirubin, direct bilirubin, alkaline phosphatase, and γ-glutamyl transpeptidase were measured to observe changes in liver function, while serum creatinine, blood urea nitrogen, and β2-microglobulin were measured to observe changes in renal function. Liver and kidney functions were observed before and 24 h after dosing. Individuals who received multiple doses of compound Danshen tablets were also monitored on day 3 (before and 24 h after dosing), day 6 (before daily dosing), and day 9 (before and 24 h after dosing).

### Preparation and NMR-based identification of major Bingpian metabolites

Individually housed rats were kept in metabolic cages, and their urine collection tubes were maintained at -15°C. Three rats were orally administered *dl*-borneol (100 mg/kg/day) for five consecutive days after being given 2.5 mL of water via gavage. Urine samples were collected at 0–8 and 8–24 h after each dose. Similarly, another set of three rats were orally given *dl*-isoborneol at 100 mg/kg/day for five consecutive days, and their urine was collected using the same method as for *dl*-borneol. The rat experiments were performed at Laboratory Animal Center of Shanghai Institute of Materia Medica (Shanghai, China). The study protocols were reviewed and approved by the Institutional Animal Care and Use Committee. The rats were treated according to the Guidance for Ethical Treatment of Laboratory Animals (the Ministry of Science and Technology of China, 2006). Male Sprague-Dawley rats were obtained from Shanghai JieSiJie Laboratory Animal (Shanghai, China). CO_2_ was used to euthanize all rats after the experiment was over.

Rat urine collected after dosing *dl*-borneol was used to isolate the Bingpian metabolite **M1**_G_. The urine was acidified with 4 M HCl and extracted twice with ethyl acetate. The combined extract was subjected to vacuum evaporation to remove the solvent. The residue was redissolved in 50% acetonitrile and subjected to HPLC separation using a Shiseido Capcell Pak C_18_ AQ 5-μm column (250 × 4.6 mm i.d.; Tokyo, Japan). The mobile phase of acetonitrile/water (30:70, v/v), containing 25 mM formic acid, was delivered isocratically at 4 mL/min. The effluent containing the target metabolite was collected and subjected to vacuum evaporation to remove the solvent. The purity of the residue was checked before dissolving in deuterated dimethyl sulfoxide. The chemical structure of the target metabolite was elucidated through ^1^H NMR, ^13^C NMR, DEPT135, ^1^H-^13^C HSQC, and ^1^H-^13^C HMBC using a Bruker AVANCE III HD 600 MHz NMR spectrometer (Bremen, Germany). The Bingpian metabolite **M2**_G_ was isolated from urine collected from *dl* -isoborneol-dosed rats and analyzed by NMR.

### Supportive in vitro studies

#### In vitro metabolism studies of Bingpian compounds

To characterize Bingpian metabolites detected in humans, NADPH-fortified human liver microsomes, NADPH-fortified human intestine microsomes, NAD-fortified human liver cytosol, NAD-fortified human intestine cytosol, UDPGA-fortified human liver microsomes, and UDPGA-fortified human intestine microsomes were incubated with borneol (final concentration, 2 µM) or isoborneol (2 µM). The detailed incubation conditions were the same as described previously [23⍰25], and sampling were done at 15 and 30 min and 1, 2, and 4 h after initiating incubation. In addition, in vitro two-step metabolism (i.e., oxidation→glucuronidation) of borneol (**1**), isoborneol (**2**), and camphor (**3**) was assessed by incubating the test compounds separately with NADPH-fortified human liver microsomes for 4 h, followed by adding UDPGA-fortified human liver microsomes and incubating for an additional 4 h. The incubation conditions were the same as described previously [26]. Prior to their use, human liver microsomes and intestine microsomes were functionally validated for their expression of P450 and UGT enzymes, respectively, by quantifying the hydroxylation of midazolam (a positive substrate of P450s) and glucuronidation of 4-methylumbelliferone (a positive substrate of UGTs). Meanwhile, human liver cytosol and intestine cytosol were assessed for alcohol dehydrogenase expression by measuring the oxidation of its positive substrate, cinnamyl alcohol.

The enzyme kinetics of the oxidation of borneol (**1**) and isoborneol (**2**) into camphor (**3**) was evaluated with respect to Michaelis constant (*K*_m_) and maximum velocity (*V*_max_) using NADPH-fortified human liver microsomes. The assessment was conducted under linear metabolism conditions via incubation for 15 min, while the test compounds’ final concentrations was within the range of 1.95 to 8000 µM. In addition, the enzyme kinetics of glucuronidation of **1** and **2** into borneol-2-*O*-glucuronide (**M1**_G_) and isoborneol-2-*O*-glucuronide (**M2**_G_), respectively, were assessed by using UDPGA-fortified human liver microsomes with resp*K*e_m_ct to and *V*_max_. The analysis was performed under linear metabolism conditions via incubation for 10 min, while the test compound concentration ranges from 1.95 to 500 µM.

To identify which human P450 isoenzymes could mediate oxidation of borneol (**1**) and isoborneol (**2**) into camphor (**3**), the cDNA-expressed human P450 enzymes, i.e., CYP1A2, CYP2A6, CYP2B6, CYP2C8, CYP2C9, CYP2C19, CYP2D6, CYP2E1, CYP3A4, and CYP3A5 (50 pmol/mL each), were fortified with NADPH and then incubated with **1** or **2** (5 μM each) for 30 min. The enzymes that could mediate the oxidation of **1** and **2** was identified based on the formation of **3**. In addition, to identify which human UGT isoenzymes could mediate glucuronidation of **1** and **2** into borneol-2-*O*-glucuronide (**M1**_G_) and isoborneol-2-*O*-glucuronide (**M2**_G_), respectively, the cDNA-expressed human UGT enzymes, i.e., UGT1A1, UGT1A3, UGT1A6, UGT1A7, UGT1A8, UGT1A9, UGT1A10, UGT2B4, UGT2B7, UGT2B10, UGT2B15, and UGT2B17 (0.25 mg protein/mL each), were each fortified with UDPGA, and then incubated with **1** or **2** (5 μM each) for 30 min. The enzymes that could mediate the glucuronidation of **1** and **2** were identified based on the formation of **M1**_G_ and **M2**_G_, respectively.

#### In vitro transport studies of Bingpian compounds

To identify the mechanism through which hepatic borneol-2-*O*-glucuronide (**M1**_G_) and isoborneol-2-*O*-glucuronide (**M2**_G_) enter systemic circulation or are excreted into bile, inside-out membrane vesicles expressing MRP3, MRP4, MRP2, BCRP, BSEP, or MDR1 were used to assess the efflux of the test compounds at 20 μM for each using a rapid filtration method described previously [27]. Before use, the membrane vesicles were functionally validated by measuring vesicular transport of oestradiol 17β-D-glucuronide (a positive substrate of MRP2, MRP3, and MRP4) and glycyrrhizin (BCRP, BSEP, and MDR1). These substrates exhibited net transport ratios of 7.8–24.6. Transport was measured in pmol/min/mg protein using the equation (1):

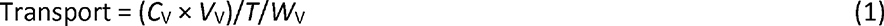

where *C*_V_ is the concentration of the compound in the vesicular lysate supernatant in μM, *V*_V_ is the volume of the lysate in μL, *T* is the incubation time (10 min), and *W*_V_ is the amount of vesicle protein amount per well. A net transport ratio (Transport_ATP_/Transport_AMP_ ratio) > 3 was considered a positive result for ATP-dependent transport. The kinetics of vesicular transport of **M1**_G_ and **M2**_G_, which are transported by human MRP3 and MRP4, were evaluated in terms of *K*_m_ and *V*_max_. The compounds were incubated under conditions of linear uptake for 5 minutes at concentrations ranging from 1.95 to 1000 μM, with the exception of **M2**_G_ transported by MRP4, which was incubated within the range of 1.95 to 500 μM. To assess the tubular secretion of circulating **M1**_G_ and **M2**_G_, transfected HEK-293 cells expressing human OAT1, OAT2, OAT3, OAT4, OATP1B1, OATP1B3, and OATP2B1 were used to assess the uptake of the test compounds (20 μM each) as previously described [28]. Before use, the transfected cells were functionally validated with estrone-3-sulfate (a positive substrate of OAT3, OAT4, and OATP2B1), para-aminohippuric acid (OAT1), prostaglandin F2α (OAT2), and oestradiol 17β-D-glucuronide (OATP1B1 and OATP1B3). These substrates exhibited net transport ratios of 27.2–184. A net transport rate ratio (Transport _TC_/Transport_MC_ ratio) > 3 was considered a positive result for differential cellular uptake mediated by a SLC transporter between the transfected cells (TC) and the mock cells (MC). The kinetics of cellular uptakes of **M1**_G_ and **M2**_G_ mediated by human OAT3 and OAT4 were assessed with respect to *K*_m_ and *V*_max_. The assessment was performed under linear uptake conditions by incubating for 5 min, and the final concentrations of the test compounds ranged from 4.7‒625 and 39‒5000 μM for OAT3 and OAT4, respectively.

#### Supportive in vitro plasma-protein-binding studies of Bingpian compounds

An equilibrium dialysis method described previously [29] was used to determine unbound fractions of borneol, isoborneol, camphor, borneol-2-*O*-glucuronide, and isoborneol-2-*O*-glucuronide in human plasma. In brief, Spectra/Por 2 dialysis membrane (molecular weight cutoff, 12–14 kDa; Rancho Dominguez, CA, USA) was used to contain plasma samples spiked with the test compounds at the final concentrations 0.2, 2, and 20 μM with incubation time 12 h at 37°C and 60 rpm to achieve equilibrium. Dialysates and associated plasma preparations were sampled for further analysis.

### Analysis of Bingpian compounds in biomatrices

Metabolite profiling of plasma and urine samples from human subjects who ingested compound Danshen tablet was conducted through gas chromatography/mass spectrometry (GC-MS) and liquid chromatography/mass spectrometry (LC-MS/MS). GC-MS analysis was conducted using Trace GC Ultra Gas Chromatograph (Thermo Fisher Scientific, Milan, Italy) connected to a DSQ II single quadrupole mass spectrometer (Thermo Fisher Scientific, Austin, TX, USA) to analyze volatile oxidized metabolites of borneol (**1**) and isoborneol (**2**), as well as the unchanged compounds. Meanwhile, LC-MS/MS analysis was performed using a Waters Synapt G2 high-definition time-of-flight mass spectrometer (Manchester, UK) with a LockSpray source and a Waters Acquity ultra-performance liquid chromatographic (UPLC) separation module (Milford, MA, USA) to analyze non-volatile conjugated metabolites. A list of metabolite candidates was generated to facilitate profiling analysis, where molecular mass gains and losses of candidate metabolites were compared with molecular masses of parent compounds **1** and **2** [30]. Sample preparation was conducted via n-hexane extraction (1:1, v/v) for plasma or urine in GC-MS-based analysis and via methanol treatment (1:3 for plasma and 1:18 for urine, v/v) in LC-MS/MS-based analysis. The chromatographic separation was achieved using a TR-5MS capillary column (30 m × 0.25 mm × 0.25 μm i.d.; Thermo Fisher Scientific, Cheshire, UK) with a helium carrier gas (99.999% purity) delivered at 1.5 mL/min through a 9-min GC gradient program (0–1 min at 60 °C, 1–7 min from 60 °C to 150 °C (at 15 °C/min), 7–8 min at 240 °C (at 90 °C/min), and 8–9 min at 240 °C). The electron impact of 70 eV was used for mass spectrometry in a selected ion monitoring mode, with transfer line and ion source both set at 250 °C. For LC-MS/MS-based analysis, chromatographic separation was achieved using a Waters Acquity UPLC HSS T3 1.8 μm column (100 × 2.1 mm i.d.; Dublin, Ireland; maintained at 45 °C) with a mobile phase comprising solvent A (water/acetonitrile, 99:1, v/v; containing 25 mM formic acid) and solvent B (water/acetonitrile, 1:99, v/v; containing 25 mM formic acid) delivered at 0.3 mL/min. A 26-min LC gradient program was followed (0–1 min at 5% solvent B, 1–20 min from 5% to 30% solvent B, 20–22 min from 30% to 55% solvent B, 22–24 min at 95% solvent B, and 24–26 min at 5% solvent B), followed by mass spectrometry in resolution mode with approximately 20000 resolving power. The electrospray ion source was operated in the negative ion mode (capillary, -2.5 kV). A scan time of 0.3 s was used for MS^E^ data acquisition (in Centroid, *m/z* 50–1500) with low collision energy (trap collision energy, 4 V) and high collision energy (ramp trap collision energy, 30–50 V).

The Bingpian circulating compounds borneol (**1**), isoborneol (**2**), and camphor (**3**) were quantified in human plasma and urine using GC-MS, while borneol-2-*O*-glucuronide (**M1**_G_) and isoborneol-2-*O*-glucuronide (**M2**_G_) were quantified using LC-MS/MS. The GC-MS-based analysis was performed on the same instrument as the metabolite profiling analysis, whereas the LC-MS/MS-based analysis utilized an Applied Biosystems SCIEX Triple Quad ^TM^ 5500 mass spectrometer (Toronto, Ontario, Canada) interfaced via a Turbo V ion source with an Agilent 1290 Infinity II LC system (Waldbronn, Germany). For GC-MS-based quantification analysis, sample preparation and chromatographic separation conditions were kept consistent with the GC-MS-based metabolite profiling analysis; MS detection utilized selected ion monitoring of **1**, **2**, **3**, and naphthalene (internal standard) at *m/z* 95, 95, 95, and 128, respectively. For LC-MS/MS-based quantification analysis, the sample preparation conditions remained unchanged from those employed in LC-MS/MS-based metabolite profiling analysis, while chromatographic separation utilized a YMC-Triart C_18_ 1.9-μm column (50 × 2.1 mm i.d.; Kyoto, Japan) and a 6-min gradient program (0–0.5 min at 20% solvent B, 0.5–4 min from 20% to 38% solvent B, 4–4.5 min from 38% to 90% solvent B, 4.5–5.0 min at 90% solvent B, 5.0–5.1 min from 90% to 20% solvent B, and 5.1–6.0 min at 20% solvent B). MS-MS detection was achieved via negative ion mode employing multiple reaction monitoring of **M1**_G_, **M2**_G_, and *l*-menthol-*O*-glucuronide (internal standard) at *m/z* 329.1→85.0 (collision energy, 36 V), 329.1→75.0 (32 V), and 331.1→75.0 (26 V), respectively. Matrix-matched calibration was attained by weighted (1/ *X*^2^) linear regression of the peak area (*Y*) versus the corresponding nominal analyte concentration (*X*, nM). Validation of assay was performed according to the Guidelines for Validation of Analytical Method Adopted in Pharmaceutical Quality Specification in the Chinese Pharmacopoeia (2020) [31]. The lower limits of quantification for **1**, **2**, **3**, **M1**_G_, and **M2**_G_ in plasma were 7.8, 7.8, 15.6, 7.8, and 7.8 nM, respectively, while in urine were 7.8, 7.8, 15.6, 78, and 78 nM, respectively. Accuracy and precision values of bioassay were 85.5–106.7% and 4.5–11.5%, respectively, for **1**, **2**, and **3**, and 101–105% and 4.02–8.08%, respectively, for **M1**_G_ and **M2**_G_. All validation results were within acceptable ranges.

The in vitro study samples were analyzed for the presence of Bingpian metabolites using the assay for the metabolite profiling. Midazolam, 4-methylumbelliferone, oestradiol 17β-D-glucuronide, glycyrrhizin, estrone-3-sulfate, para-aminohippuric acid, prostaglandin F2α, and oestradiol 17β-D-glucuronide were analyzed using previously reported assays [23, 27]. Furthermore, the concentration of borneol (**1**), isoborneol (**2**), camphor (**3**), borneol-2-*O*-glucuronide (**M1**_G_), and isoborneol-2-*O*-glucuronide (**M2**_G_) in the in vitro study samples was quantified using assays designed for measuring these compounds in human samples.

### In vitro assessment of effect of major circulating Bingpian compounds on ox-LDL-induced macrophage foaming

Based on the results of the preceding pharmacokinetic study of Bingpian in humans who orally received compound Danshen tablets, five major circulating Bingpian compounds, i.e., borneol-2-*O*-glucuronide (**M1**_G_), isoborneol-2-*O*-glucuronide (**M2**_G_), camphor (**3**), borneol (**1**), and isoborneol (**2**), were assessed in vitro for their ability to inhibit ox-LDL-induced macrophage foaming. Atorvastatin was used as a positive control [32–34]. The test Bingpian and control compounds were dissolved initially in DMSO and then diluted with DMEM, which was supplemented with 1% FBS and 1% penicillin-streptomycin (with a DMSO concentration ≤ 0.1% for final use).

#### Cytotoxicity assay

Cytotoxicity potential of the test Bingpian compounds against RAW264.7 cells was assess using CCK-8. The cells were incubated in a 5% CO_2_ humidified incubator at 37°C for 12 h in DMEM supplemented with 10% FBS and 1% penicillin-streptomycin. Following treatment with atorvastatin (5, 10, and 20 μM), the test Bingpian compounds (20, 100, and 500 μM per compound concentration), and **MBC** (a mixture of these test Bingpian compounds at concentrations similar to their respective unbound maximum plasma concentration after dosing compound Danshen tablets in humans, at a total concentration of 13.2 μM) for 24 h in DMEM that was supplemented with 1% FBS and 1% penicillin-streptomycin, the cells (in 100 μL of medium per well) were further treated with ox-LDL (100 μg/mL) for 24 h, then treated with CCK-8 reagent (10 μL) for an additional 111h. DMEM (with a final DMSO concentration of 0.1%) served as a negative control. The absorbance of the incubation was measured at 450 nm using a Molecular Devices SpectraMax M2 plate reader (Sunnyvale, CA, USA). The survival percentage was calculated by normalizing to the negative control.

#### Foam cell staining

The Oil Red O (ORO) stain kit was used, following the manufacturer’s instructions, to visualize lipid droplets in ox-LDL-treated macrophages. RAW264.7 cells were subjected to treatment with test Bingpian compounds (20 μM each), **MBC**, and atorvastatin (5 μM) for 24 h in DMEM supplemented with 1% FBS and 1% penicillin-streptomycin after overnight cell culture, then with ox-LDL (100 μg/mL) for another 24 h. Upon incubation completion, cells were washed twice with PBS followed by fixation in 4% paraformaldehyde for 30 min. Cells were then stained with ORO at 37 °C for 1 h for lipid droplets and, then with hematoxylin for 1 min for cell nuclei. Cell images were acquired using a Nikon ECLIPSE Ti-S fluorescence microscope (Tokyo, Japan), and the ORO-stained area was estimated using Image-Pro Plus software (version 6.0; Media Cybernetics, Rockville, MD, USA). Foam cells were scored if they contained ≥ 10 lipid droplets[35].

#### Cholesterol measurement

To quantify the intracellular levels of cholesterol after treating macrophages subjected to ox-LDL treatment, the total and free cholesterol assay kit were employed as per the provided guidelines. Overnight cultured RAW264.7 cells were treated with the test Bingpian compounds (20 μM/screen), **MBC**, and atorvastatin (5 μM) individually for 24 h in 1% FBS and 1% penicillin-streptomycin supplemented DMEM, followed by the addition of ox-LDL (100 μg/mL) for another 24 h. Further, cells were washed twice with PBS and subjected to cell lysis for 10 min. After centrifugation of the lysate, supernatant was mixed with the reagents R1 and R2 (at a ratio of 4:1, v/v) and incubated at 37 °C for 20 min. Cholesterol ester was estimated by subtracting free cholesterol from total cholesterol. Normalization of all data was done with respect to the protein level of the corresponding samples, using the bicinchoninic acid protein assay kit, in alignment with the provided instruction.

#### DiI-ox-LDL uptake assay

To evaluate the potential inhibition of ox-LDL uptake into macrophages by the test Bingpian compounds, DiI-ox-LDL was used for the assessment. After cell culture overnight, RAW264.7 cells were treated separately with atorvastatin (5 μM), the test Bingpian compounds (20 μM each), and **MBC** for 24 h in DMEM supplemented with 1% FBS and 1% penicillin-streptomycin and then with DiI-ox-LDL (50 μg/mL) for an additional 4 h. Following the incubation period, the cells were washed thrice with PBS and fixed in 4% paraformaldehyde for 30 min, and stained for cell nuclei with DAPI for 4 min. Cell images were acquired on the Nikon ECLIPSE Ti-S fluorescence microscope and the mean fluorescence intensity was estimated using Image-Pro Plus software.

#### 22-NBD-cholesterol efflux assay

To evaluate whether the test Bingpian compounds promote cholesterol efflux from macrophages, 22-NBD-cholesterol was employed for the assessment. After being cultured for 4 h, RAW264.7 cells were initially incubated with ox-LDL (100 μg/mL) and 22-NBD-cholesterol (5 µg/mL) for 14 h in phenol-free DMEM supplemented with 1% FBS and 1% penicillin-streptomycin following which the cells were washed twice with PBS. The cells were subsequently treated with the cholesterol acceptor HDL (50 µg/mL), atorvastatin (5 μM), the test Bingpian compounds (20 μM each), and **MBC** separately for an additional 4 h, after which, the supernatant and cell lysate were collected. To measure fluorescence intensity, we used the Molecular Devices SpectraMax M2 plate reader, with excitation and emission wavelengths of 469 nm and 538 nm, respectively. The cholesterol efflux rate was calculated by dividing the intensity of supernatant by total intensity of supernatant and lysate.

### Data processing

Plasma pharmacokinetic parameters of Bingpian compounds were estimated by non-compartmental analysis using Kinetica software (version 5.0; Thermo Scientific, Philadelphia, PA, USA). GraFit software (version 5; Surrey, United Kingdom) was used to determine *K*_m_ and *V*_max_ via nonlinear regression analysis of initial transport rates as a function of substrate concentration. All data are expressed as the mean ± standard deviation. Statistical analysis was conducted in GraphPad Prism software (version 8.3.0; San Diego, CA, USA), and a *P* value < 0.05 was considered statistically significant.

## RESULTS

### Metabolites are the predominant circulating compounds of Bingpian in humans

Like in the national drug reference standard sample of synthetic Bingpian, borneol (**1**) and isoborneol (**2**) were the two major constituents of the component Bingpian in compound Danshen tablet (GuoYaoZhunZi-Z53021243), while camphor (3) was a minor Bingpian constituent. In the current human pharmacokinetic study, the compound doses of **1**, **2**, and **3** from the tablet were 462, 329, and 1.1 μmol/person, respectively. No significant adverse event was reported during the period of the human study. Furthermore, dosing the tablet did not induce any abnormal change in liver or kidney function indicators. After dosing the tablet, **1**, **2**, and **3** were detected in human plasma by GC - MS, but at very low level sinurine (Fig. 1; Table 1). Inconsistent with their compound doses, the level of systemic exposure to **3** (in area under the plasma concentration-time curve from zero to 48 h after dosing/AUC _0-48h_) was significantly higher than those of **1** and **2**. This suggests that the vast majority of circulating **3** was a metabolite rather than absorbed as the unchanged compound. In addition, 18 other Bingpian metabolites were also detected in plasma and/or urine by LC-MS/MS (Fig. 1; Table 2); they were not detected in compound Danshen tablet. Two major glucuronidated metabolites demonstrated significantly elevated levels of systemic exposure compared to their parent compounds **1** and **2**, and were predominantly excreted into urine. Meanwhile, the other metabolites detected in human samples were oxidized-glucuronidated variants with low exposure levels.

**Fig. 1.**
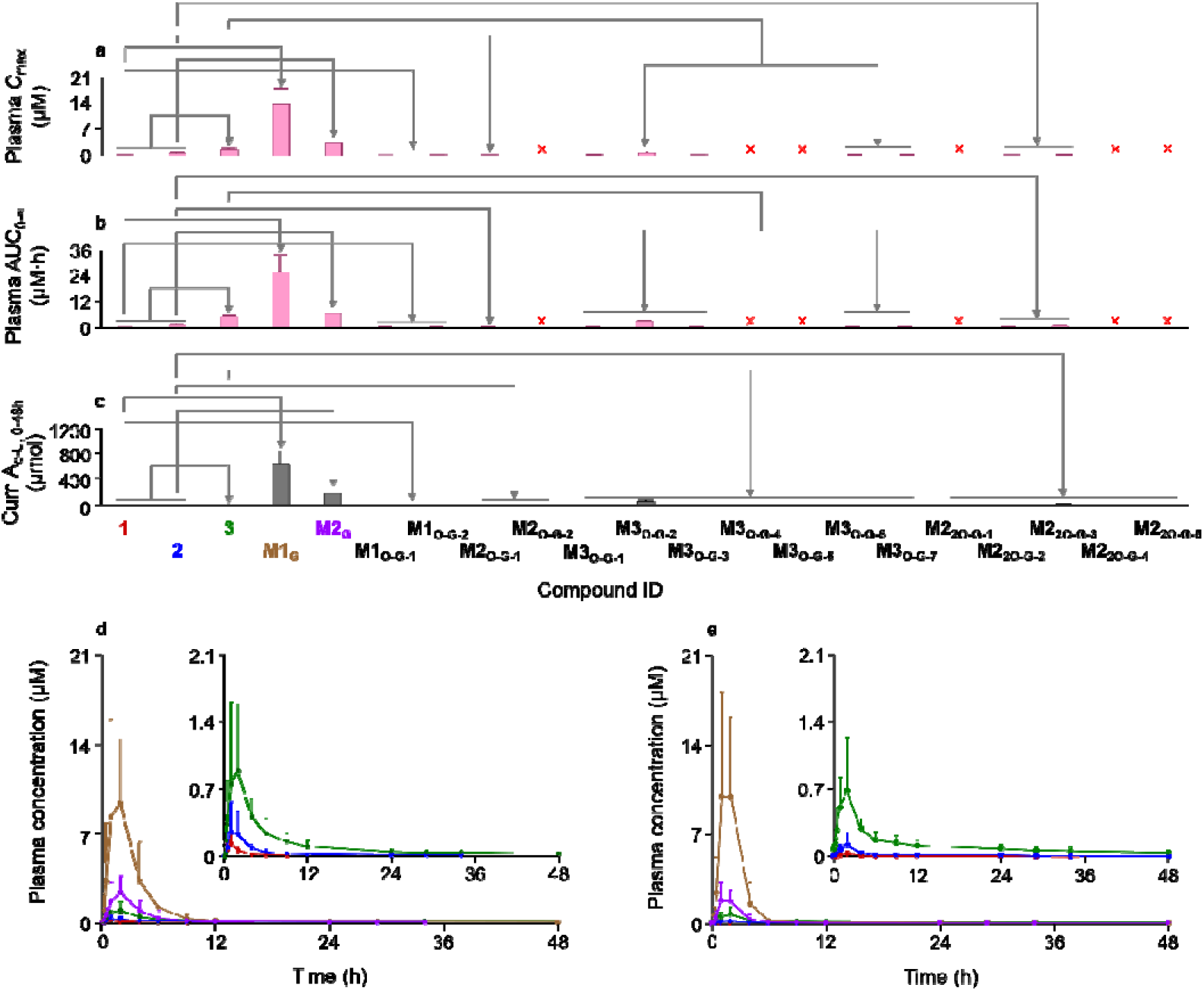
Systemic exposure to and metabolism of Bingpian compounds in human subjects after dosing compound Danshen tablet, a Bingpian-containing Chinese herbal medicine. **a** systemic exposure data; **b** renal excretion data; **c** and **d** mean plasma concentration-time profiles of **1** (red line), **2** (blue line), camphor (**3**; green line), borneol-2-*O*-glucuronide (**M1**_G_; brown line), and isoborneol-2-*O*-glucuronide (**M2**_G_; purple line) before and after dosing the tablet on day 1 and day 9, respectively. The human pharmacokinetic study is described in the **MATERIALS AND METHODS** section. The chemical names of other Bingpian metabolites, shown as their compound ID, are listed in Table 2. The red symbol “×” denotes the compound that was not detected.

**Table 1.** Bingpian compounds detected, using GC-MS, in plasma and urine of human subjects who were given Compound Danshen tablets.

| ID | Compound name | GC-MS data |  | Molecular formula | Molecular mass (Da) | Occurrence in human samples |
| --- | --- | --- | --- | --- | --- | --- |
| | | $t_R$ (min) | Ionization profile<sup>a</sup> ($m/z$) | | | |
| **1** | Borneol | 6.24 | **95**, 110, 121, 154 | C<sub>10</sub>H<sub>18</sub>O | 154.1358 | Plasma, urine |
| **2** | Isoborneol | 6.39 | **95**, 110, 121, 154 | C<sub>10</sub>H<sub>18</sub>O | 154.1358 | Plasma, urine |
| **3** | Camphor | 6.47 | **95**, 81, 108, 152 | C<sub>10</sub>H<sub>16</sub>O | 152.1201 | Plasma, urine |

<sup>a</sup> Number in bold, product ion of base peak.

**Table 2.**
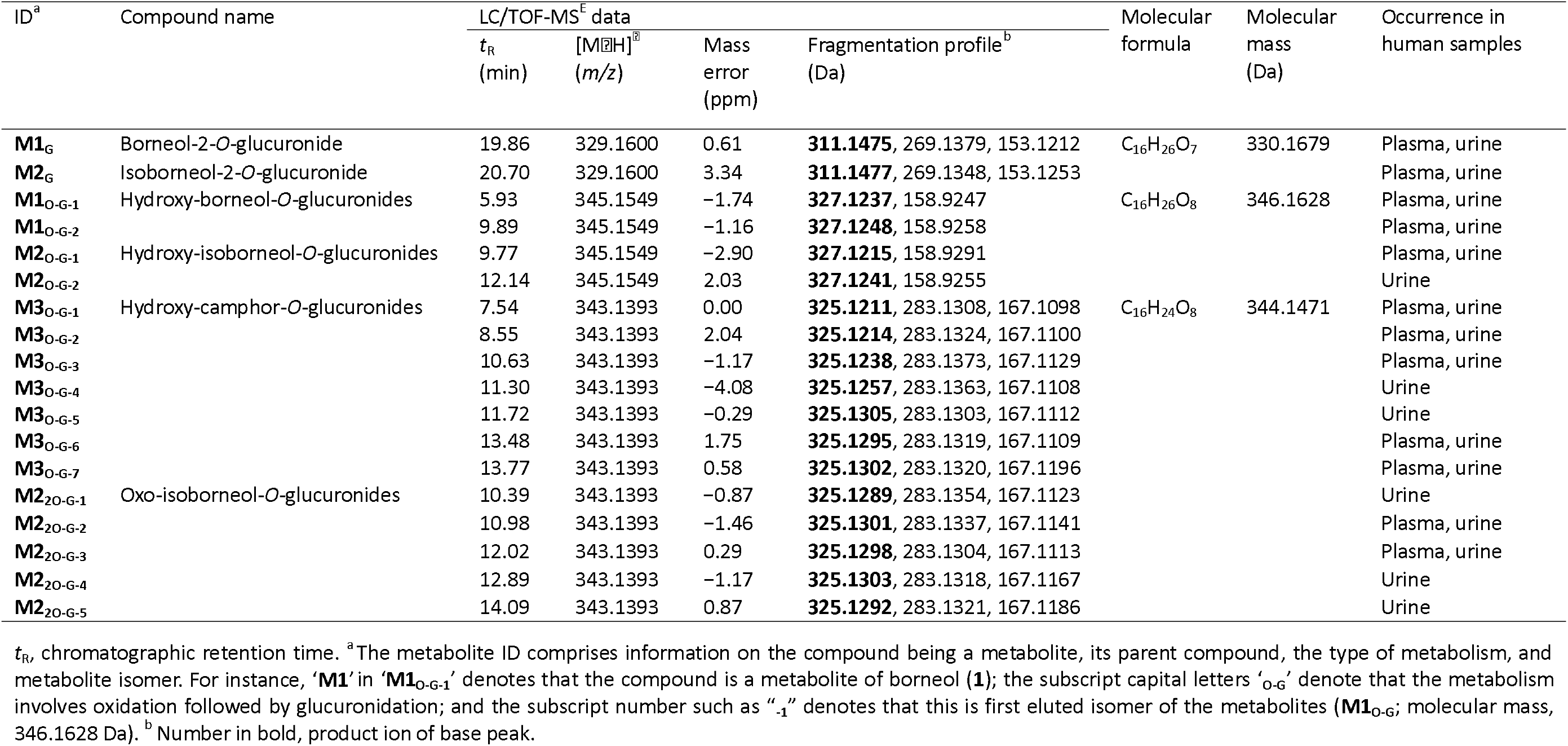
Bingpian compounds detected, using LC-MS/MS, in plasma and urine of human subjects who were given Compound Danshen tablets.

To identify the parent compounds of the two major glucuronides, a rat pharmacokinetic study was conducted by separately administering borneol (**1**) and isoborneol (**2**) as they are isomers. Consistent with the situation in humans, the two glucuronidated metabolites were separately the major circulating metabolite of **1** and that of **2** in rats and both extensively excreted into rat urine. In rats, **1** produced camphor (**3**) and one of the two glucuronides, while **2** produced 3 as well and the other glucuronides. Both the glucuronides were extensively excreted into rat urine. These two metabolites were isolated from rat urine collected after repeatedly dosing individual compound **1** or **2**. The isolated metabolites were analyzed using NMR and LC-MS/MS and characterized as borneol-2-*O*-glucuronide (**M1**_G_) and isoborneol-2-*O*-glucuronide (**M2**_G_), which are the metabolites of compounds 1 and 2 respectively (see Table 3).

**Table 3.**
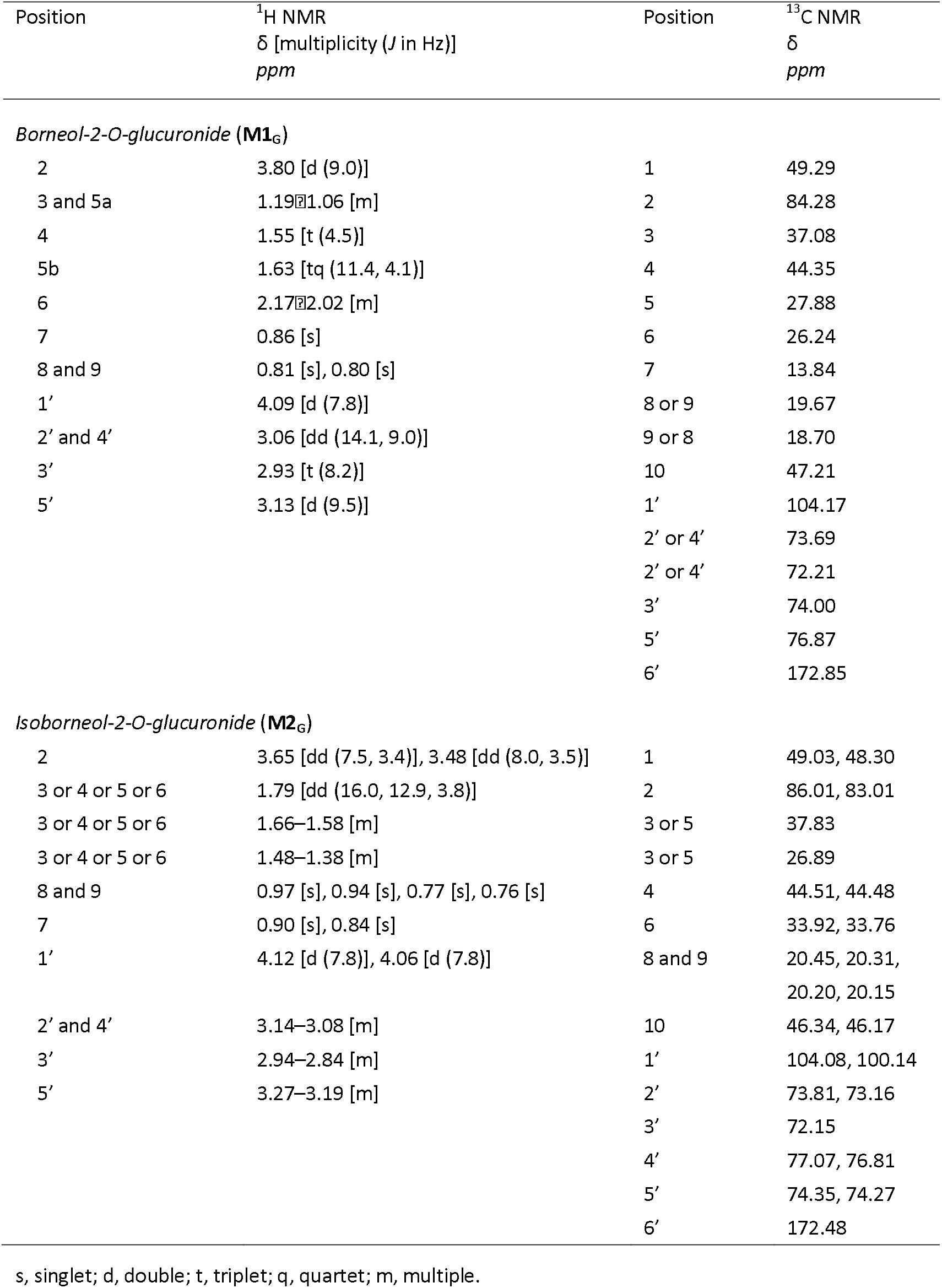
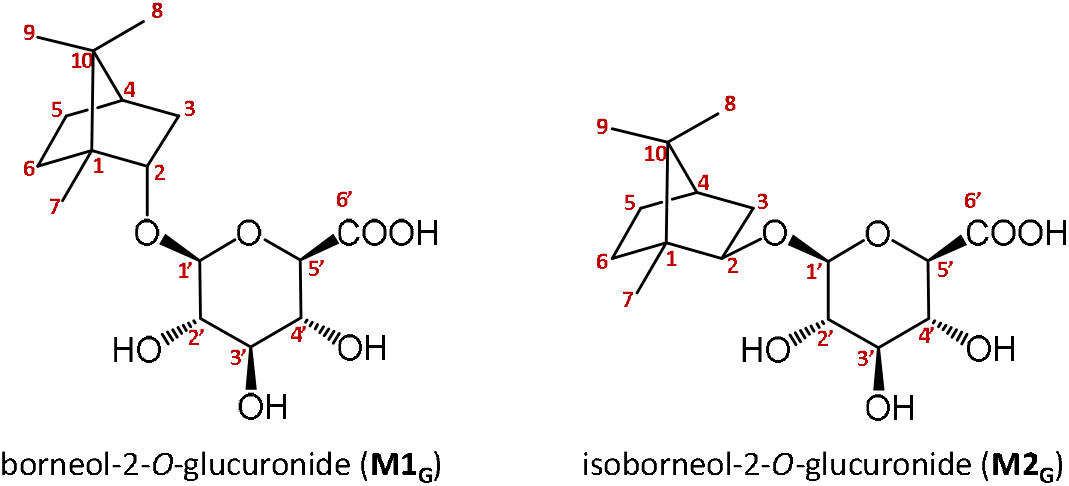
NMR data of borneol-2-*O*-glucuronide (**M1_G_**) and isoborneol-2-*O*-glucuronide (**M2_G_**).

Table 4 summarizes the pharmacokinetic parameters of the circulating Bingpian compounds borneol (**1**), isoborneol (**2**), camphor (**3**), borneol-2-*O*-glucuronide (**M1**_G_), and isoborneol-2-*O*-glucuronide (**M2**_G_) in human subjects who orally received compound Danshen tablet, with the plasma concentration-time profiles illustrated in Fig. 1. The plasma concentrations of the five compounds showed a similar pattern after administration, with increases and decreases occurring in a parallel manner. This was indicated by their inter-compound similarity in the peak concentration (*T*_peak_), half-life (*t*_1/2_), and mean residence time (MRT). Renal excretion of the parent compounds **1** and **2** and the metabolite **3** was poor, but that of **M1**_G_ and **M2**_G_ was extensive. The renal clearance ratios [CL _R_/(GFR×*f*_u_)] showed that renal excretion of the parent compounds and the metabolite involved tubular reabsorption, while that of the glucuronides involved tubular secretion. The results indicated that the glucuronidation process enhanced elimination of **1** and **2** through the renal excretion of their glucuronides. Furthermore, there was a significant difference in *f*_e-U_ between **M1**_G_ and **M2**_G_, which seemed to be dose-dependent based on compound Danshen tablet (refer to Table 4 note). It is worth pointing out that in vivo inter-conversions between **M1**_G_ and **M2**_G_, as well as between **1** and **2**, were not observed when rats were administered with pure **1** and **2** separately. No significant gender difference was found among the human subjects for the five Bingpian circulating compounds in *C*_max_, AUC _0-48h_, *t*_1/2_, or MRT (*P* = 0. 2–41.00; after adjusting the compound doses for the respective subjects’ body weights, except for *t*_1/2_ or MRT). Repeatedly dosing compound Danshen tablet in human subjects for seven consecutive days did not result in any significant accumulation of the circulating compounds (Fig. 1).

**Table 4.** Pharmacokinetic parameters of circulating Bingpian compounds after administering the compound Danshen tablets to human subjects.

| Pharmacokinetic parameter | Circulating Bingpian compound (ID) |  |  |  |  |
| --- | --- | --- | --- | --- | --- |
|  | Borneol (1) | Isoborneol (2) | Camphor (3) | Borneol-2-*O*-glucuronide (M1<sub>G</sub>) | Isoborneol-2-*O*-glucuronide (M2<sub>G</sub>) |
| *Plasma data* |  |  |  |  |  |
| $C_{\max}$ (μM) | 0.11 ± 0.09 | 0.36 ± 0.28 | 1.27 ± 0.73 | 13.7 ± 4.54 | 3.13 ± 0.89 |
| $T_{\text{peak}}$ (h) | 1.9 ± 1.3 | 2.1 ± 1.2 | 2.4 ± 1.5 | 2.0 ± 1.0 | 2.3 ± 1.6 |
| $AUC_{0-48\text{h}}$ (h·μM) | 0.18 ± 0.13 | 1.11 ± 0.67 | 5.35 ± 2.28 | 29.1 ± 6.11 | 7.31 ± 1.64 |
| $t_{1/2-(1)}$ (h) | 1.4 ± 0.9 | 2.4 ± 0.8 | 3.1 ± 0.5 | 1.4 ± 0.1 | 1.6 ± 0.2 |
| $t_{1/2-(2)}$ (h) | — | 13.1 ± 4.4 | 18.1 ± 7.1 | 9.1 ± 1.9 | 7.5 ± 1.6 |
| MRT (h) | 3.1 ± 1.9 | 8.2 ± 3.7 | 11.3 ± 3.6 | 3.5 ± 1.2 | 3.9 ± 1.1 |
| *Urine data* |  |  |  |  |  |
| $Cum.A_{e-U}$ (μmol) | 0.05 ± 0.03 | 0.02 ± 0.01 | 0.2 ± 0.2 | 607 ± 191 | 173 ± 59 |
| $f_{e-U}$ (%) | 0.01 ± 0.01 | 0.005 ± 0.005 | 0.03 ± 0.03 | 131 ± 41* | 53 ± 18* |
| $CL_R/(GFR \times f_u)$ | 0.22 ± 0.25 | 0.01 ± 0.005 | 0.02 ± 0.02 | 4.7 ± 2.0 | 5.7 ± 2.2 |

$C_{\max}$ , maximum plasma concentration; $T_{\text{peak}}$ , the time taken to achieve the peak plasma concentration; $AUC_{0-48\text{h}}$ , area under the plasma concentration-time curve from zero to 48 h after dosing; $t_{1/2-(1)}$ , first half-life; $t_{1/2-(2)}$ , second half-life; MRT, mean residence time; $Cum.A_{e-U}$ , cumulative amount excreted into urine from zero to 48 h; $f_{e-U}$ , fractional urinary excretion; $CL_R$ , renal clearance; GFR, glomerular filtration rate; $f_u$ , unbound fraction in plasma. \*In a pilot study dosing human subjects with compound Danshen tablet at 9 tablets/person, $f_{e-U}$ of borneol-2-*O*-glucuronide (M1<sub>G</sub>) and isoborneol-2-*O*-glucuronide (M2<sub>G</sub>) was 107 ± 1.3% and 88.7 ± 11.6%, respectively.

### Multiple hepatic enzymes and transporters could influence systemic exposure to metabolites of borneol and isoborneol

To verify the Bingpian metabolites detected in human plasma and urine, the catalytic capability of human hepatic enzymes was compared in vitro with that of intestinal enzymes. Human liver microsomes fortified with NADPH had significant catalytic capability for oxidation of borneol (**1**) and isoborneol (**2**) into camphor (**3**), while the catalytic capability of human intestine microsomes was poor. Additionally, only NAD-fortified human liver cytosol, but not the corresponding cytosol from human intestine, could mediate the oxidation of *d*-borneol to **3**, rather than of *l*-borneol and isoborneol, and was inhibited by 4-methylpyrazole (a specific inhibitor of alcohol dehydrogenase). For glucuronidation of **1** and **2** into the corresponding glucuronides, UDPGA-fortified human liver microsomes had significant ability to catalyze, while the catalytic capability of human intestine microsomes was poor. Furthermore, incubation of **3** and NADPH-fortified human liver microsomes, followed by the addition of UDPGA, produced seven metabolites: **M3**_O-G-1_, **M3**_O-G-2_, **M3**_O-G-3_, **M3**_O-G-4_, **M3**_O-G-5_, **M3**_O-G-6_, and **M3**_O-G-7_. Incubation of 1 and NADPH-fortified human liver microsomes, followed by the addition of UDPGA, yielded the metabolites **M1**_O-G-1_, **M1**_O-G-2_, and the seven metabolites of **M3**_O-G_; such metabolic reactions were poor with human intestine microsomes. Similar scenario took place for **2**, and the detected metabolites were **M2**_O-G-1_, **M2**_O-G-2_, **M2**_2O-G-1_, **M2**_2O-G-2_, **M2**_2O-G-3_, **M2**_2O-G-4_, **M2**_2O-G-5_, and the seven metabolites of M3_O-G_. These metabolic reactions of human intestine microsomes were poor. Collectively, the liver is the major metabolic site for **1** and **2**. No metabolic conversion was observed between **1** and **2** or between **M1**_G_ and **M2**_G_ in in vitro metabolism studies.

The classic Michaelis-Menten equation best fits the UGT-mediated glucuronidation, while biphasic kinetics was observed for the P450-mediated oxidation. Figure 2a and b shows representative plots of metabolite formation rate versus substrate concentrations for the metabolic turnover of borneol (**1**) and isoborneol (**2**). Glucuronidation of **1** into borneol-2-*O*-glucuronide (**M1**_G_) was mediated primarily by UGT2B7, while glucuronidation of **2** into isoborneol-2-*O*-glucuronide (**M2**_G_) was exclusively by UGT2B7 (Fig. 2c). Likewise, oxidation of **1** into camphor (**3**) was mediated primarily by CYP2A6, CYP3A4, and CYP3A5, while a similar scenario occurred for the oxidation of **2** into **3**, except for CYP2B6 also playing a major role (Fig. 2d). Based on the results of the metabolite profiling of human samples from the in vivo pharmacokinetic study and the in vitro metabolism studies, we proposed pathways for the metabolism of **1** and **2** in humans (Fig. 3).

**Fig. 2.**
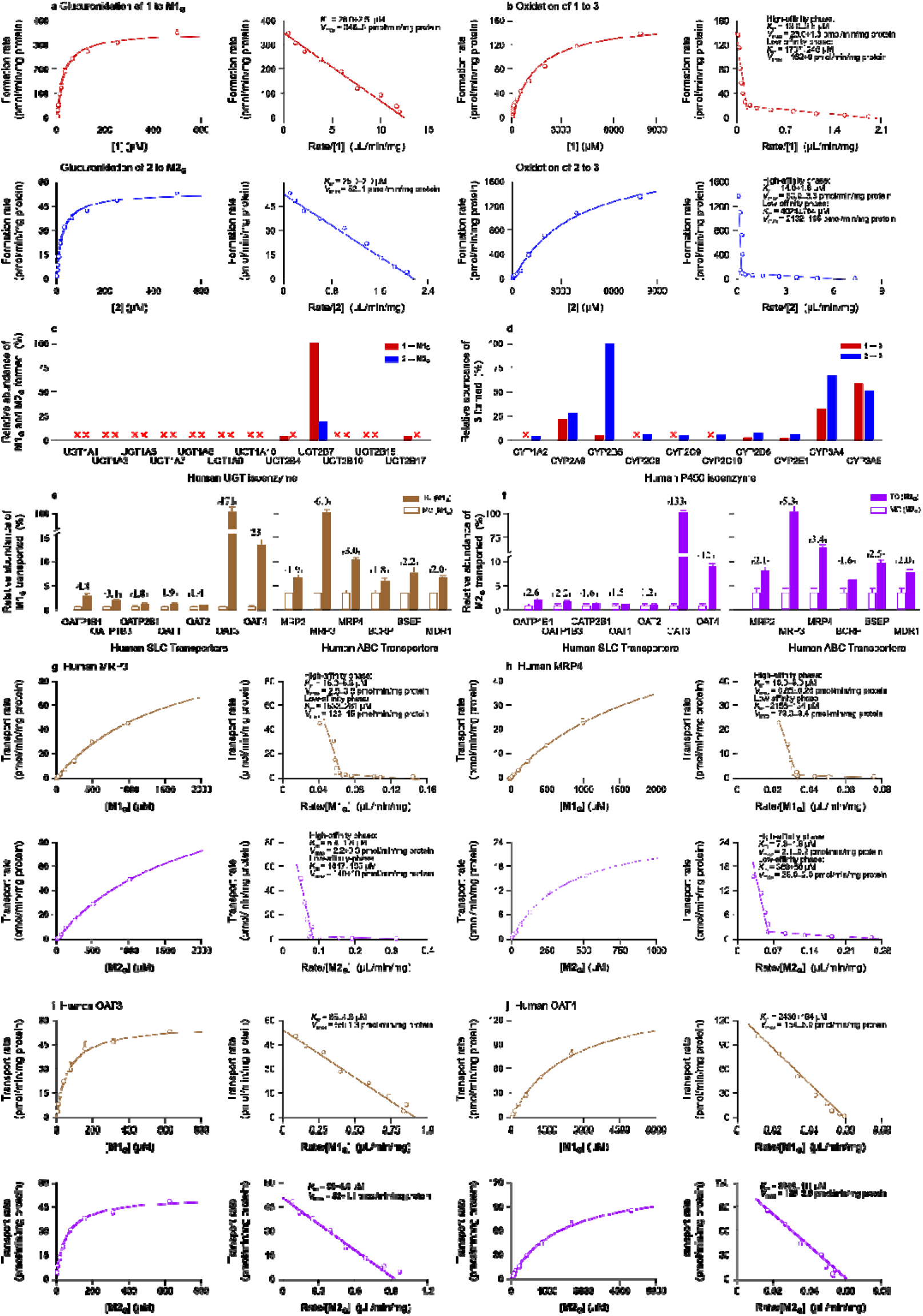
Enzyme kinetics of glucuronidation and isoenzymes responsible for of borneol (1) and isoborneol (2) and for oxidation of the compounds, as well as transporters and associated kinetics of transport vs substrate concentration for transport of borneol-2-*O*-glucuronide (M1_G_) and isoborneol-2-*O*-glucuronide (M2_G_). **a** the kinetic plot of formation of M1_G_ and M2_G_ vs concentration of **1** and **2**, respectively, and the associated Eadie-Hofstee plots; **b** the kinetic plot of formation of camphor (**3**) vs concentration of **1** and **2**, respectively, and the corresponding Eadie-Hofstee plots; **c** the glucuronidation activities of cDNA-expressed human UGT isoenzymes for the substrates **1** and **2**; **d** the oxidation activities of cDNA-expressed human P450 isoenzymes for the substrates **1** and **2**; **e** and **f** net transport ratios for in vitro transport of **M1_G_** and **M2_G_** mediated by human hepatic and renal transporters, respectively. **g** and **h** representative kinetic plots of transport vs substrate concentration for the efflux of **M1_G_**and **M2_G_** mediated by human MRP3 and MRP4 and cellular uptake of **M1_G_** and **M 2_G_** mediated by human OAT3 and OAT4. *K*_m_, Michaelis constant; *V*_max_, maximum velocity.

**Fig. 3.**
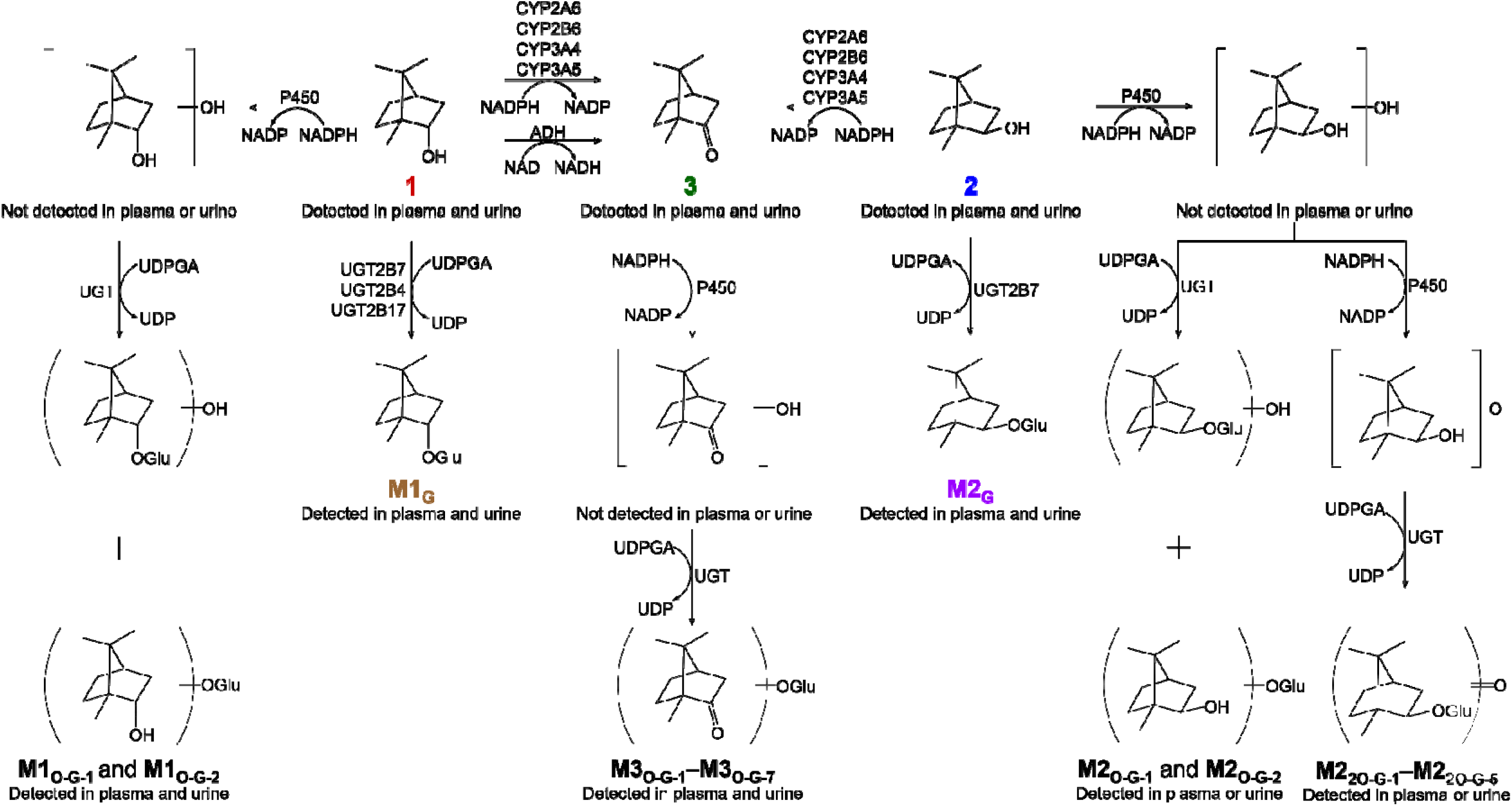
Proposed major metabolic pathways of the Bingpian compounds borneol (1) and isoborneol (2). The chemical names of Bingpian metabolites, shown as their compound ID, are listed in Table 1 and 2. ADH, alcohol dehydrogenase; CYP and P450, cytochrome P450 enzyme; Glu, glucuronosyl; NAD, nicotinamide adenine dinucleotide; NADP, β-nicotinamide adenine dinucleotide phosphate; NADPH, reduced β-nicotinamide adenine dinucleotide phosphate; UDP, uridine 5’-diphosphate; U DPGA, uridine 5’-diphosphoglucuronic acid; UGT, uridine 5’-diphosphoglucur onosyltransferase.

As shown inFig. 2e and f, in vitro vesicular transport study suggested that both borneol-2-*O*-glucuronide (**M1**_G_) and isoborneol-2-*O*-glucuronide (**M2**_G_) were substrates of human hepatic MRP3 and MRP4; this results in the efflux of these glucuronides into bloodstream following their hepatic formation. Conversely, neither **M1**_G_ nor **M2**_G_ were found to be substrates for human hepatic MRP2, BCRP, BSEP, or MDR1, indicating that the hepatobiliary excretion of the glucuronides was limited. Furthermore, **M1**_G_ and **M2**_G_ were also substrates of human renal OAT3 and OAT4, which initiate the tubular secretion of the glucuronides. Figure 2g–j shows representative plots of transport versus substrate concentration for the vesicle transport and cellular uptake of **M1**_G_ and **M2**_G_. Unbound fractions (*f*_u_) in human plasma were 41.0%, 31.5%, 56.9%, 62.6%, and 63.7% for borneol (**1**), isoborneol (**2**), camphor (**3**), **M1**_G_, and **M2**_G_, respectively.

### Bingpian metabolites reduce ox-LDL-induced lipid accumulation and foam-cell formation in RAW264.7 cells

Based on the results of the preceding human pharmacokinetic study, five circulating Bingpian compounds, i.e., borneol (**1**), isoborneol (**2**), camphor (**3**), borneol-2-*O*-glucuronide (**M1**_G_), and isoborneol-2-*O*-glucuronide (**M2**_G_), were tested in vitro for their inhibition of ox-LDL-induced lipid accumulation and foam-cell formation in macrophages. The CCK-8 assay indicated that cellular toxicity of the Bingpian compounds was low (toxic concentration, ≥ 500 μM; Fig. 4a). In contrast, atorvastatin (positive control in the anti-atherosclerotic studies) exhibited high cellular toxicity with a toxic concentration of 10 μM. Ox-LDL treatment (100 μg/mL) induced lipid accumulation and foam-cell formation in RAW264.7 cells, as shown in Fig. 4b and c. Co-treatment with atorvastatin (5 μM and for all in vitro studies except the CCK-8 study) significantly attenuated the effect (*P* <0.01). The test Bingpian compounds **1**, **2**, **3**, **M1**_G_, and **M2**_G_ (20 μM each and for all the in vitro studies except the CCK-8 study) considerably inhibited lipid accumulation and foam-cell formation (*P* <0.01). Furthermore, **MBC** at a total concentration of 13.2 μM, which was estimated as a sum of the individual unbound *C*_max_ values of the five Bingpian compounds after dosing compound Danshen tablets in human subjects, also significantly inhibited foam-cell formation (*P* <0.01). Similarly, ox-LDL treatment significantly increased intracellular cholesterol ester levels in RAW264.7 cells (100 μg/mL; *P* <0.05), but pretreatment with atorvastatin significantly reduced such levels compared to ox-LDL treatment alon*P*e <(0.01; Fig. 4d). Moreover, pretreatment with **1**, **2**, **3**, **M1**_G_, **M2**_G_, and MBC separately also significantly reduced intracellular levels of cholesterol ester compared to ox-LDL treatment alone (*P* <0.01 for all).

**Fig. 4.**
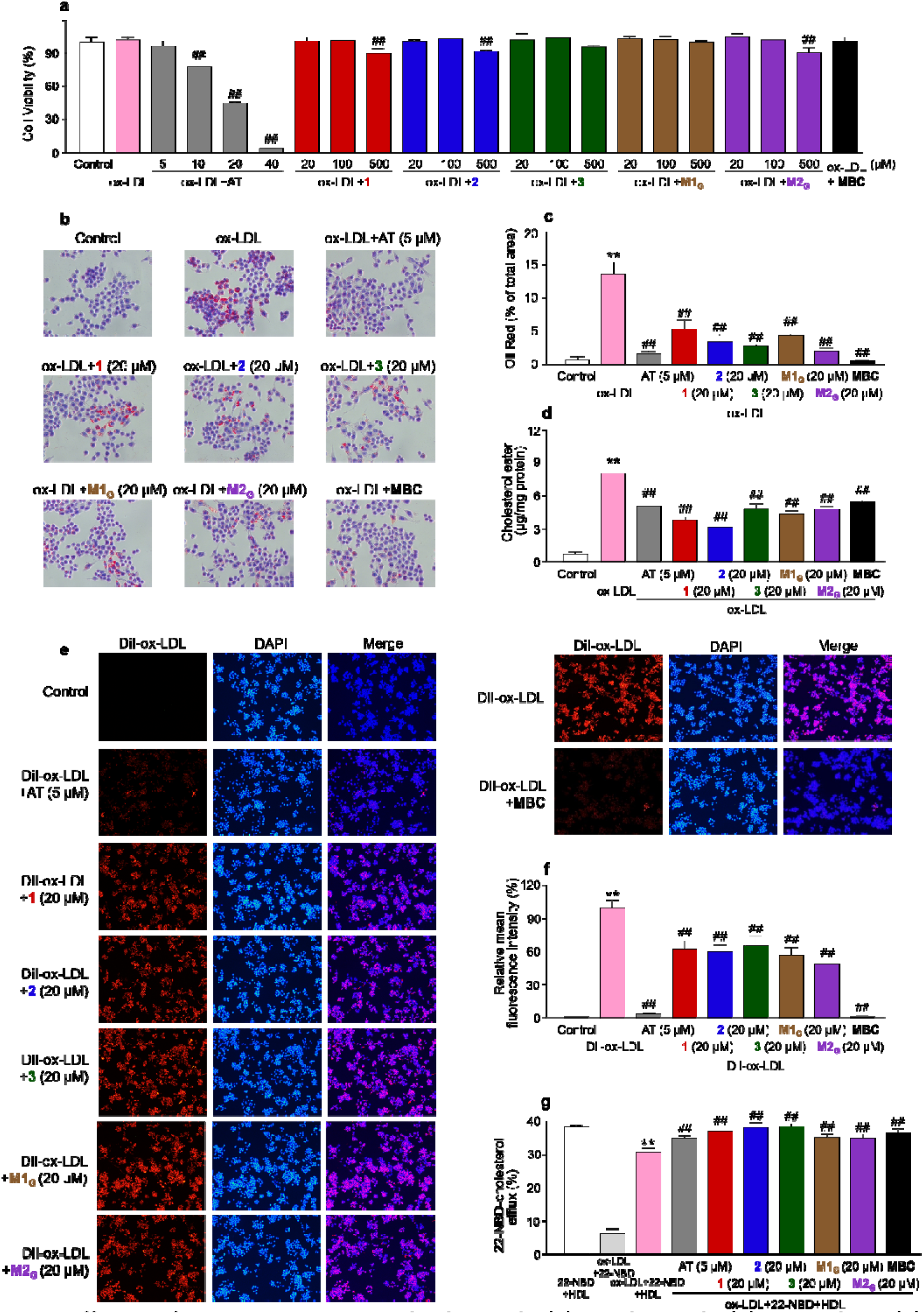
Effects of Bingpian compounds borneol (1), isoborneol (2), camphor (3), borneol-2-*O*-glucuronide (M1_G_), isoborneol-2-*O*-glucuronide (M2_G_), and their combination MBC on macrophage foaming induced by ox-LDL. **a** the cytotoxicity potential of the test Bingpian compounds and MBC against RAW264.7 cells; **b** and **c** representative images (magnified at X400) and quantitative analysis of Oil Red O-stained RAW264.7 cells treated with the test Bingpian compounds and **MBC**; **d** normalized cholesterol ester levels based on cellular protein content in RAW 264.7 cells; **e** and **f** representative images (magnified at X100) and quantitative data of DiI-ox-LDL uptake assay for the test Bingpian compounds and **MBC** in RAW264.7 cells; **g** quantitative data of 22-NBD-cholesterol efflux assay for the test Bingpian compounds and **MBC**. \*\**P* < 0.01 versus vehicle control; ^##^*P* < 0.01 versus model control.

**Fig. 5.**
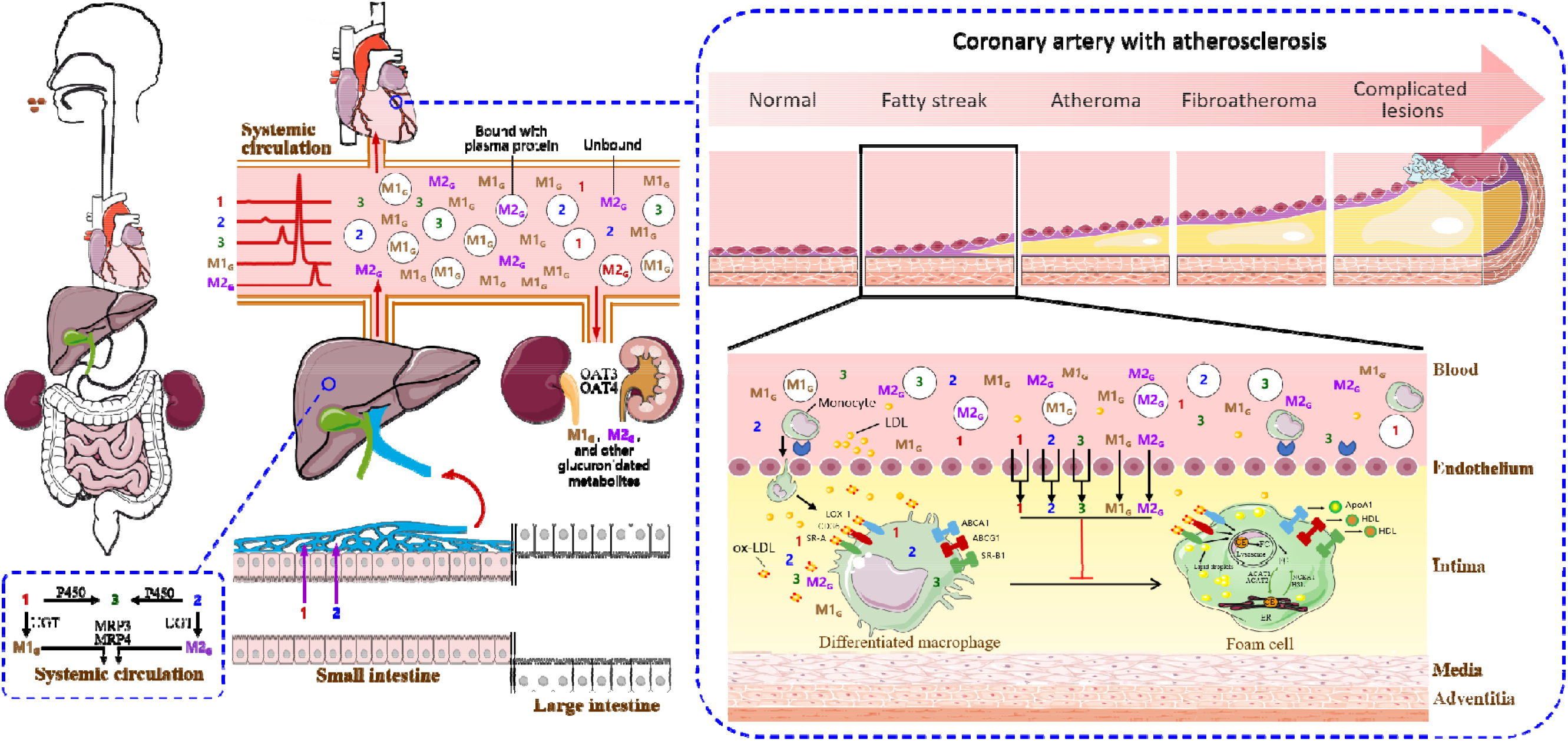
A schematic overview of formation and disposition of circulating Bingpian metabolites that inhibit foam-cell formationnmiacrophages induced by ox-LDL. The Bingpian constituents, i.e., borneol (**1**) and isoborneol (**2**), are efficiently absorbed in the intestine. However, after extensive hepatic first-pass glucuronidation (mainly mediated by UGT2B7, coupled with MRP3 and MRP4-mediated efflux of the glucuronides into the blood) and oxidation (media ted by CYP2A6, CYP2B6, and CYP3A), the metabolites borneol-2-O-glucuronide (**M1_G_**), isoborneol-2-O-glucuronide (**M2_G_**), and camphor (**3**), rather than the unchanged **1** and **2**, become the major circulating Bingpian compounds in the systemic circulation. **M1_G_** and **M2_G_** undergo renal excretion involving both glomerular filtration and tubular secretion mediated by OAT3 and OAT4, while **3** is eliminated via glucuronidation followed by renal excretion. Early stages of atherosclerosis trigger various events such as endothelial injury, activation, and dysfunction, culminating in reduced endothelial barrier function. The glucuronides **M1_G_** and **M2_G_**, unlike **1**, **2**, and **3**, exhibit low membrane permeability. Because the endothelial monolayer has diminished, **M1_G_** and **M2_G_**can cross the bloodstream and reach macrophages at the arterial intima. Fatty streaks, the early visible atherosclerotic lesions, mainly comprise macrophage foam cells that substantially take up cholesterol, leading to plaque buildup in the arteries. The Bingpian compounds **M1_G_**, **M2_G_**, **3**, **1**, and **2** inhibit ox-LDL-induced lipid accumulation and foam-cell formation in macrophages and could act synergistically. Given the levels of systemic exposure and inhibitory activities on foam-cell formation in macrophages, the metabolites must be given priority in pharmacodynamic studies of Bingpian. The clipart used in this figure has been adapted from Servier’s Medical Art database (https://smart.servier.com). ACAT1 and ACAT2, cholesterol acyltransferase-1 and -2, respectively; CE, cholesteryl esters; ER, endoplasmic reticulum; FC, free cholesterol; HSL, hormonesensitive lipase; NCEH1, neutral cholesteryl ester hydrolases 1.

To better understand if the observed inhibition of lipid accumulation and foam-cell formation in macrophages is primarily due to decreased uptake of ox-LDL, increased efflux of cholesterol, or a combination of both, the DiI-ox-LDL uptake assay and cholesterol efflux assay were performed. As shown in Fig. 4e and f, fluorescence intensity of RAW264.7 cells significantly amplified upon treatment with DiI-ox-LDL (50 μg/mL; *P* <0.01), but such intensity was significantly reduced when the cells were pretreated with atorvastatin, in contrast to the intensity by DiI-ox-LDL-treatment alone (*P* <0.01). Additionally, separate pretreatment with borneol (**1**), isoborneol (**2**), camphor (**3**), borneol-2-*O*-glucuronide (**M1**_G_), isoborneol-2-*O*-glucuronide (**M2**_G_), and **MBC** resulted in a substantial decrease in fluorescence intensity (*P* <0.01 for all). Consequently, the test Bingpian compounds exhibited the potential to hinder macrophage uptake of ox-LDL. Fig. 4g demonstrates that atorvastatin substantially amplified the efflux of 22-NBD-cholesterol from RAW264.7 cells that were exposed to ox-LDL, compared to the ones without atorvastatin-treatment (*P* <0.01). Furthermore, separate treatment with **1**, **2**, **3**, **M1**_G_, **M2**_G_, and MBC resulted in a significant boost in the cellular efflux of 22-NBD-cholesterol (*P* <0.01 for all). Therefore, the test Bingpian compounds demonstrated the potential to encourage cellular efflux of cholesterol. Collectively, both inhibiting cellular uptake of ox-LDL and promoting cellular efflux of cholesterol are responsible for Bingpian’s attenuation of ox-LDL-induced lipid accumulation and foam-cell formation in macrophages by its major metabolites and the parent compounds.

## Discussion

Accurate and translational pharmacological research on a Chinese herbal medicine requires the pharmacodynamic studies that focused on ‘right’ compounds of the medicine. This lays the foundation for precisely understanding how the herbal medicine functions in the body to deliver therapeutic benefits. To this end, pharmacokinetic studies are conducted to single out the potentially important compounds (accessible to the action site with significant levels of therapeutically relevant body exposure) of the medicine for the pharmacodynamic studies [36, 37]. Such potentially important compounds can not only be the parent compounds but the metabolites as well. Prior pharmacodynamic studies were conducted only with parent borneol and isoborneol [16–18]. However, our current pharmacokinetic study, by dosing the Bingpian-containing herbal medicine compound Danshen tablet in humans, demonstrated that the levels of systemic exposure to the metabolites borneol-2-*O*-glucuronide (**M1**_G_), isoborneol-2-*O*-glucuronide (**M2**_G_), and camphor (**3**) were significantly higher than the levels of the parent compounds borneol (**1**) and isoborneol (**2**). As the parent compounds and the metabolites have similar in vitro potency of inhibiting lipid accumulation and foam-cell formation in macrophages (via decreased uptake of ox-LDL and increased efflux of cholesterol), researchers should be cautious when attempting to relate plasma concentrations of **1** and **2** alone to the effect of Bingpian following oral administration. Instead, the circulating metabolites **M1**_G_, **M2**_G_, and **3** should be of great therapeutic concern in the pharmacological research on Bingpian or Bingpian-containing medicines.

Glucuronides are commonly considered to be inactive or minimally active metabolites of drugs. While this holds true for most drugs, there are some exceptions [38, 39]. Morphine-6-*O*-glucuronide, for instance, shares the anesthetic property with the parent drug morphine and increases its effect [40]. In addition, ezetimibe-*O*-glucuronide has lipid-lowering property along with the parent drug ezetimibe, augmenting the effect [41], while darexaban *-O*-glucuronide has antithrombotic property along with darexaban, augmenting the effect [42]. Our current investigation offers another instance of bioactive glucuronides, derived from Bingpian. Besides their pharmacodynamic activities as a prerequisite for exerting pharmacological effects in vivo, glucuronides must meet two pharmacokinetic prerequisites: significant level of exposure and accessibility to the action sites. After oral administration of compound Danshen tablet, significant levels of systemic exposure to borneol-2-*O*-glucuronide (**M1**_G_) and isoborneol-2-*O*-glucuronide (**M2**_G_) occur due to rapid hepatic glucuronidation of borneol (**1**) and isoborneol (**2**), predominantly by UGT2B7, followed by efflux of the glucuronides from the hepatocytes into the blood by MRP3 and MRP4. Unlike the nonpolar parent compounds **1** and **2**, the polar glucuronides **M1**_G_ and **M2**_G_ have low membrane permeability and are eliminated from the systemic circulation exclusively by renal excretion (involving glomerular filtration and OAT3/4-mediated tubular secretion). The second pharmacokinetic prerequisite for the in vivo inhibition of lipid accumulation and foam-cell formation in macrophages by **M1**_G_ and **M2**_G_ is their accessibility to macrophages present in the arterial intima. During the development of atherosclerosis, endothelial injury, activation, or dysfunction is an early event, which reduces the endothelial barrier function [43, 44]. The transfer of **M1**_G_ and **M2**_G_ across the impaired endothelial monolayer, from the bloodstream to the arterial intima, allows these glucuronides to reach macrophages.

Multiple scavenger receptors on the cell membrane of differentiated macrophages, such as CD36, SR-A, and LOX-1, uptake atherogenically modified LDL (such as ox-LDL) in response to endothelial injury [45]. The major circulating Bingpian compounds can be divided into two types: borneol-2-*O*-glucuronide (**M1**_G_) and isoborneol-2-*O*-glucuronide (**M2**_G_), which have poor membrane permeability and thus can only access the scavenger receptors extracellularly, and camphor (**3**), borneol (**1**), and isoborneol (**2**), which have good membrane permeability and can access the receptors both extracellularly and intracellularly. Although both types of Bingpian compounds (20 μM for each) exhibit inhibitory activities on uptake of ox-LDL, MBC (a mixture of **M1**_G_, **M2**_G_, **1**, **2**, and **3** at a total concentration of 13.2 μM, relevant to the therapy of compound Danshen tablet) exhibited significantly more potent activity than the individual compounds, suggesting that there are synergistic effects among the compounds. Whether the synergistic effects of the Bingpian compounds on ox-LDL uptake result from extracellular action and intracellular downregulation of the scavenger receptors remains investigation. Similar scenario is expected to take place in promoting efflux of cholesterol by these compounds. Although cytosolic β-glucuronidase present in macrophages is capable of deglucuronidating **M1**_G_ and **M2**_G_ into **1** and **2**, respectively, poor membrane permeability restricts the cellular entry of the glucuronides to access the enzyme. The recovery of 92% of **M1**_G_ and **M2**_G_, with the limited detection of **1** and of **2** in the medium after exposing RAW264.7 cells to the glucuronides, indicates that the glucuronides inhibit lipid accumulation and foam cell formation in macrophages independently, rather than through their deglucuronidated products.

Cholesterol belongs to a class of lipid molecules in the body critical for various physiological functions such as maintaining and constructing cell membranes, producing hormones, increasing nutrient absorption, and stimulating bile formation. The cholesterol cycle is a fundamental process that maintains lipid homeostasis by balancing the synthesis, utilization, and elimination of cholesterol from the liver and other organs [46]. Fatty streaks, the earliest visible atherosclerotic lesions, are primarily made of macrophage foam cells that have taken up massive amounts of cholesterol [47]. Over time, the foam cells may result in plaque buildup in the arteries, posing a significant risk in atherosclerosis progression. Inhibition of foam-cell formation is emerging as an attractive strategy for therapeutic intervention of atherosclerosis in clinics [45, 47]. Strategies to inhibit foam-cell formation in macrophages involve multiple approaches such as reducing lipid uptake mediated by multiple scavenger receptors, decreasing cholesterol esterification by acetyl-CoA acetyltransferases, and promoting cholesterol efflux mediated by ABCA1 transporter (ApoA-1 as the acceptor), ABCG1 (HDL as the acceptor), and SR-B1 (HDL also as the acceptor) [45]. Our investigation shows that multiple Bingpian compounds circulating in the bloodstream, particularly the metabolites borneol-2-O-glucuronide (**M1**_G_), isoborneol-2-*O*-glucuronide (**M2**_G_), and camphor (**3**), act synergistically to inhibit ox-LDL-induced lipid accumulation and foam-cell formation in macrophages. As the anti-atherosclerotic action of Bingpian is multifaceted, it is important to investigate how these compounds interact with the preceding receptors, enzymes, and transporters in macrophages and modulate associated pathways. Specifically, it is necessary to determine if Bingpian compounds act independently on different targets within the network or if each compound acts on multiple targets. Additional strategies to reduce foam-cell formation include decreasing LDL modification and inhibition of monocyte to macrophage differentiation [45]. Further investigation of the Bingpian compounds is necessary to determine their effects on foam-cell formation via these approaches. In addition, endothelial dysfunction increases the risk of foam-cell formation by facilitating lipid accumulation in the vessel wall, while foam-cell formation triggers an inflammatory response that worsens endothelial damage, creating a dangerous feedback loop that can lead to the development and progression of atherosclerosis and associated complications [43]. Recently, we found that the Bingpian compounds circulating in the bloodstream protect endothelial cells stimulated by ox-LDL (publication pending).

Integrating pharmacokinetic and pharmacodynamic studies comprehensively enables the precise and translational refinement of pharmacological research on Chinese herbal medicines, focusing on the ‘right’ compounds. In summary, the major metabolites borneol-2-*O*-glucuronide (**M1**_G_), isoborneol-2-*O*-glucuronide (**M2**_G_), and camphor (**3**) are significantly circulating compounds of Bingpian, rather than the parent compounds borneol (**1**) and isoborneol (**2**), after administering the Bingpian-containing herbal medicine compound Danshen tablet to humans. These Bingpian metabolites act synergistically to inhibit ox-LDL-induced lipid accumulation and foam-cell formation in macrophages. Future investigation of Bingpian’s anti-atherosclerotic effects should account for the divergent cellular distribution of glucuronides **M1**_G_ and **M2**_G_ (with limited penetration through the cell membrane and extracellular activity only) compared to the oxidized metabolite **3** (capable of intracellular and extracellular activity due to good membrane permeability). Furthermore, since endothelia participate in and suffer from foam-cell formation, endothelial protection of Bingpian warrants more investigation. The current investigation demonstrates the significance of the circulating metabolites in pharmacological research of Bingpian or Bingpian-containing herbal remedies.

## ACKNOWLEDGMENTS

This work was funded by grants from the National Natural Science Foundation of China (81673582), Innovation Team and Talents Cultivation Program of National Administration of Traditional Chinese Medicine (ZYYCXTD-C-202009), and the Strategic Priority Research Program of the Chinese Academy of Sciences (XDA12050306).

## AUTHOR CONTRIBUTIONS

Participated in the research design: CL, and CC; Conducted the experiments: CC, RRH, HL, ZXC, FQW, FFD, FX, JQW, and TW; Performed the data analysis: CC, CL, RRH, HL, and ZXC; Wrote or contributed to the writing of the manuscript: CL, CC, RRH, HL, ZXC, and OEO.

